# A Machine Learning and Benchmarking Approach for Molecular Formula Assignment of Ultra High-Resolution Mass Spectrometry Data from Complex Mixtures

**DOI:** 10.64898/2026.02.17.706479

**Authors:** Bilal Shabbir, Pablo R. B. Oliveira, Francisco Fernandez-Lima, Fahad Saeed

## Abstract

A machine learning approach to molecular formula assignment is crucial for unlocking the full potential of ultra-high resolution mass spectrometry (UHRMS) when analyzing complex mixtures. By combining data-driven models with rigorous benchmarking, the accuracy, consistency, and speed in identifying plausible molecular formulas from vast spectral datasets can be improved. Compared with traditional *de novo* methods that rely heavily on rule-based heuristics, and manual parameter tuning, machine learning approaches can capture complex patterns in data and adapt more readily to diverse sample types. In this paper, we describe the application of a machine learning methods using the k-nearest neighbors (KNN) algorithm trained on curated chemical formula datasets of UHRMS analysis of dissolved organic matter (DOM) covering the saline river continuum and tropical wet/dry season variability. The influence of the mass accuracy (training set with 0.15-1ppm) was evaluated on a blind test set of DOMs of different geographical origins. A Decision Tree Regressor (DTR) and Random Forest Regressor (RFR) based on mass accuracy (<1ppm) was used. Results from our ML models exhibit 43% more formulas annotated than traditional methods (5796 vs 4047), Model-Synthetic achieved 99.9% assignment rate and annotated/assigned 2x more formulas (8,268 vs 4047). DTR and RFR achieved formula-level accuracies (FA) of 86.5% and 60.4%, respectively. Overall, results show an increase in formula assignment when compared with traditional methods. This ultimately enables more reliable characterization of complex natural and engineered systems, supporting advances in fields such as environmental science, metabolomics, and petroleomics. Furthermore, the novel data set produced for this study is made publicly available, establishing an initial benchmark for molecular formula assignment in UHRMS using machine learning. The dataset and code are publicly available at: https://github.com/pcdslab/dom-formula-assignment-using-ml

**CCS CONCEPTS:** Computing methodologies → Machine Learning → Learning paradigms → Supervised Learning

## 1 Introduction

Organic matter is extraordinarily complex, comprising thousands of chemically distinct compounds. These compounds vary in structure, reactivity, and origin. Understanding this complexity is fundamental to advancing knowledge of global biogeochemical cycles and ecosystem functioning. Dissolved organic matter (DOM) is a complex mixture of organic molecules and plays a central role in water carbon cycles and aquatic ecosystems [1]. The fulvic acid fraction (FA-DOM) represents a smaller, oxidized, and soluble subset of DOM. FA-DOM is important because it is very mobile in aquatic systems and bioavailable to micro-organisms, and affects both carbon reactivity and element rotation in river [2]. Therefore, it is essential to characterize FA-DOM at the molecular level to understand how chemical composition reflects different transformations in different rivers. Ultra-high-resolution mass spectrometry (UHRMS) such as Fourier transformation ion cyclotron mass spectrometry (FT-ICR MS) [3] can detect thousands of molecular features in a single sample, and provide the mass accuracy required to assign molecular formulas to observed peaks [4]. However, a single m/z peak can correspond to multiple potential formulas within the narrow mass error window, and the high compositional diversity of FA-DOM makes the formula allocation process a major computational research challenge [5].

Traditionally, formula assignment is applied based on predefined chemical rules and element ratio constraints, including molecular indices such as hydrogen-to-carbon (H/C), oxygen-to-carbon (O/C) ratios, and double bond equivalents [6], [7]. While these rule-based algorithms [8] are widely used, they often struggle with complex mixtures [9] because of non-standard elemental combinations and environmental variability, as they can violate pre-computed constraints [10]. Additionally, when comparing complex mixture samples from rivers with different organic matter sources, these methods can yield inconsistent formula distributions, and limits the ability to make cross-system comparisons [11]. The decreased accuracy of traditional techniques, and the growing size of FT-ICR MS datasets make this a problem that can be solved using machine-learning approaches. Machine learning (ML) methods can learn the complex relationships between mass spectral features, and molecular formulas directly from data [12], [13], [14], [15]. In recent years, ML models have been used in different mass spectrometry applications such as peak classification, isotopic pattern recognition, and spectral annotation [16], [17], [18], [19]. However, the use of ML for direct molecular formula assignment in environmental samples, particularly in the context of FA-DOM, remains understudied. A major barrier is the absence of publicly available, high mass resolutions and high mass accuracy datasets which can support robust model training and evaluation.

In this paper, we propose a machine learning framework for molecular formula assignment in FA-DOM that leverages ultra-high resolution mass spectrometry FT-ICR MS data. The framework learns relationships between molecular formulas and spectral features derived from high-resolution mass-to-charge (m/z) data. We developed K-Nearest Neighbors (KNN), Decision Tree Regressor (DTR), and Random Forest Regressor (RFR) models to predict molecular formulas of FA-DOM for different river systems. KNN uses ultra-high-resolution m/z detection and m/z assignment error to learn the relationships between known chemical formulas and their m/z features. It predicts the most likely formula for unknown peaks by comparing them to their closest neighbors in the training dataset. Absence of publicly available, diverse and high-quality benchmark is one of the bottlenecks for ML models in this domain. To address this limitation, we produced ultra-high resolution mass spectrometry dataset, acquired at 7 Tesla (7T), 9.4 Tesla (9.4T) and 21 Tesla (21T) magnetic fields with mass resolution/mass accuracy of 1 PPM (L1), 0.2-0.4 PPM (L2) and 0.15 PPM (L3) respectively. Furthermore, we produced a large-scale synthetic dataset of chemically plausible CHONS molecular formulas. This dataset allows applying ML models to FA-DOM to enhance molecular formula assignment in complex environmental samples. KNN was trained on 4 different variations of data: Model-L1, Model-L3, Model-L1-L3 (Ensemble), and Model-Synthetic (Ensemble trained using L1-L3 and synthetic data set). Model-Synthetic achieved 99.9% assignment rate and Model L1-L3 annotated/assigned 43% more formulas (5,796 vs 4,047) compared to conventional methods. DTR and RFR achieved formula-level accuracies (FA) of 86.5% and 60.4%, respectively.

The contribution of this papers as follows: (1) publicly providing a ultra-high resolution FT-ICR MS dataset for multiple mass resolution (1 PPM, 0.2-0.4 PPM, 0.15 PPM) suitable for training, validating and testing of new ML models; (2) generating a large-scale synthetic dataset of chemically plausible CHONS molecular formulas; (3) training machine learning models (KNN, Decision Tree, Random Forest) on different sets/resolutions of data; (4) using ensemble learning to enhance KNN models with synthetic data.

## 2 Methods

The overall strategy is illustrated in Fig 1. We developed a K-Nearest Neighbors (KNN) based pipeline that contained four models: Model-L1, Model-L3, Model-L1-L3 (Ensemble) and Model-Synthetic (Ensemble). Each model is trained on a different set of training data to assign formulas for the L2 test set. After predicting a peak from each model, predictions were selected based on the lowest parts-per-million (ppm) error, with peaks exceeding 1 ppm being considered false annotations. To evaluate performance, we systematically varied hyperparameters (k = 1, 3 and distance metric = Euclidean, Manhattan), resulting in 16 total configurations across the four models. Furthermore, we trained Decision Tree and Random Forest models as regression problem to predict the counts of the following elements: Carbon (C), Hydrogen (H), Oxygen (O), Nitrogen (N) and Sulfur (S). We then compute the element-level accuracy (EA) and formula-level accuracy (FA) to validate these findings.

**Figure 1:**
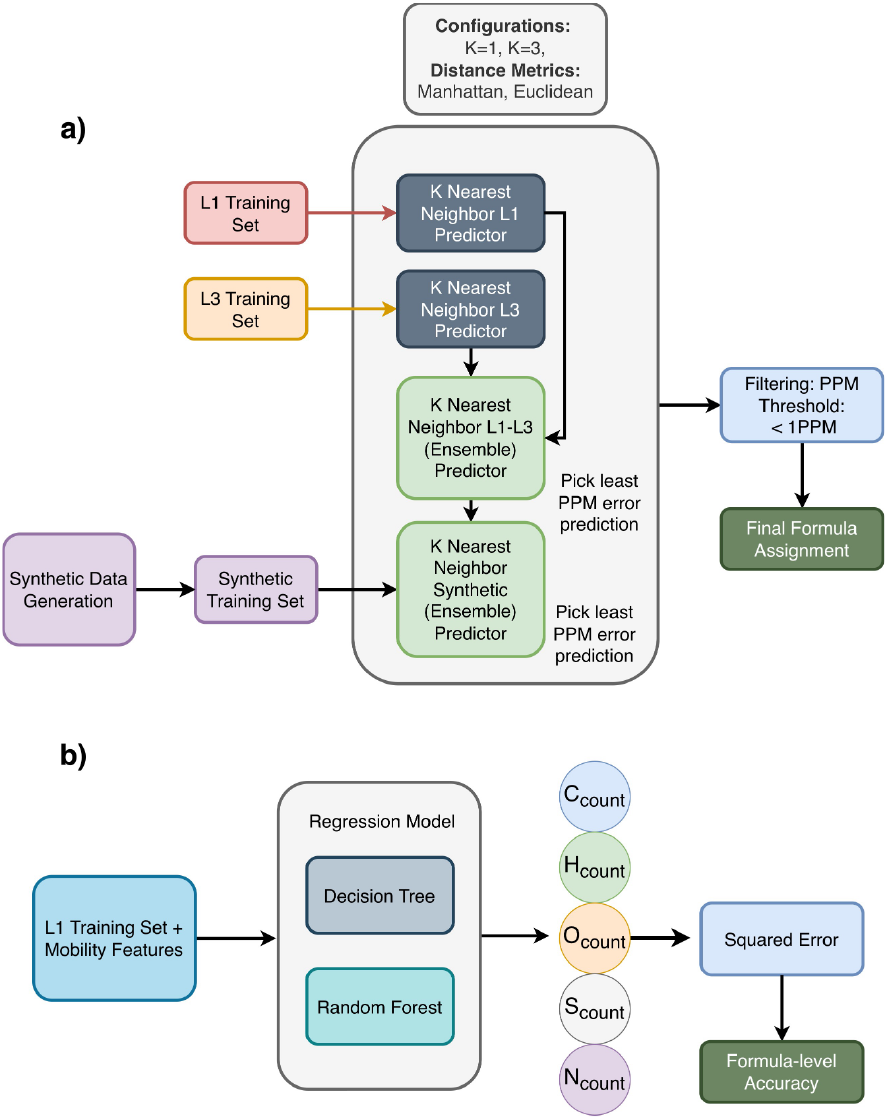
(a) KNN Pipeline for Formula Assignment/Annotation. For each train set, first KNN model is fitted, and then formula assignment/annotation is performed using the closest peaks in the training set. After formula assignment/annotation, all the predictions with more than 1 PPM error are considered false annotations. (b) Decision Tree Regressor (DTR) and Random Forest Regressor (RFR) are trained to predict element counts for CHONS using squared-error criterion.

### 2.1 Novel Datasets Generated for this study

The training set was composed of eight environmental samples from three different river systems: five from the Harney River (HR-1 to HR-5), collected along a marsh-to-estuary transect in Everglades National Park, USA, two from Pantanal National Park (PAN7 and PAN11) in southeastern Brazil and one from the Suwannee River in the Okefenokee Swamp in south Georgia - Suwannee River Fulvic Acid Standard II (SRFA2). To capture a range of data quality and complexity, all these samples were analyzed on a 7T (L1) and on a 9.4 T (L2) ESI (-) FT-ICR MS. The samples were diluted to a final concentration of 1 ppm in methanol for analysis [20]. A subset of these samples (SRFA2 and HR-1 to HR-5) was also measured on a 21T (L3) ESI-FT-ICR MS at a higher concentration of 100 ppm [21]. This multiple-instrument approach allowed the model to be trained on data with different levels of mass resolution and accuracy. The first set contained the data with mass resolution of 1 PPM (L1) and the second set contained the mass resolution 0.15 PPM (L3). For L1 training set, we also captured mobility features, which served as the training basis for DTR and RFR models.

The test set, which was kept separate from the training data, consisted of three dissolved organic matter (DOM) standards: Suwannee River Fulvic Acid Standard II (SRFA2), Suwannee River Fulvic Acid Standard III (SRFA3), and Pahokee Peat Fulvic Acid Standard III (PPFA). These standards were acquired from the International Humic Substances Society (St. Paul, MN). Each was dissolved in Optima LC-MS grade methanol. This set contained the mass resolution of 0.2-0.4 PPM (L2). We used three variations of the test set. The first variation (L2) is a subset of the above mentioned standards that contains a labeled test set in which each peak is assigned a formula using the Composer tool. The second variation (L2-peaks) is the unlabeled peak lists for which new assignments are performed. The third variation (L2-v2) is the subset of L2 set with formulas that lie in the distribution of L1 training set with mobility features. KNN was evaluated on L2 and L2-peaks. To validate DTR and RFR, we used L2-v2 of the test set that contains mobility features in addition to mass features.

For the L1 and L3 data, formula assignments were performed with DOM theoretical constraints [22], using Composer software. All spectral data were stored in a structured manner, with separate folders for each instrument’s resolution, ensuring that the test data and its corresponding reference formula tables were not included in the model training process.

### 2.2 Data Processing and Synthetic Data Generation

For both the training and test datasets, the m/z values were rounded to five decimal places to ensure consistency across sets. Duplicated *m/z* and formula pairs were removed from the training dataset.

In addition, a synthetic dataset was generated to increase the training set of the KNN model for molecular formula assignment. We created this dataset using a combinatorial approach, in which we systematically generated every possible chemical formula from five key elements: Carbon, Hydrogen, Oxygen, Nitrogen, and Sulfur. This method allowed for the creation of a theoretical dataset containing plausible molecules within a set of constraints. The elemental ranges (C4-C50, H4-H100, O1-O25, N0-N3, S0-S2) were defined based on the known properties to ensure that the generated molecules were chemically relevant [23].

For each chemical formula, the exact theoretical mass was calculated and rounded to five decimal places. The generated formulas were then filtered based on a mass range of 100.0 to 650.0 Daltons, which corresponds to the typical mass-to-charge window of interest for DOM. The formulas were filtered on the basis of oxygen vs carbon (O/C) ration and hydrogen vs carbon (H/C) ratio. O/C ratio was considered between 0 and 1, while H/C ratio was considered between 0.3 and 2.5. Double Bond Equivalents (DBE) range was in −10 and 10. The final validated formulas and their corresponding masses were organized into a clean and comprehensive dataset.

### 2.3 Approach

Our approach involved developing machine learning frameworks for DOM data. We developed K-Nearest Neighbors (KNN) pipeline using scikit-learn’s KNeighborsClassifier (Python 3.8+) with the following components. The Model-L1 was trained on the L1 training dataset and predicted formulas using the nearest neighbor peaks from this dataset. Similarly, the Model-L3 was trained on the L3 dataset. The Model-L1-L3 (Ensemble) was an ensemble of these two models. For each test set, it generated predictions from both models and selected the formula with the smallest ppm error. Model-Synthetic (Ensemble) was another ensemble that combined the outputs of the Model-L1-L3 (Ensemble) and synthetic data, keeping the prediction with the lowest ppm error. After prediction, all peaks with a ppm error of less than 1 were considered valid, while those with errors greater than 1 were false annotations. This process is illustrated in Figure 1(a).

In addition to using different datasets, we explored multiple hyperparameter configurations to evaluate the performance of the KNN-based models. Specifically, we varied the k-value (1 and 3) and the distance metric (Euclidean and Manhattan) for each predictor named, Model-L1, Model-L3, Model-L1-L3 (Ensemble), and Model-Synthetic (Ensemble). Each of these four models was trained and tested under all possible combinations of the two k-values and two distance metrics.

Since there were four models in total (Model-L1, Model-L3, Model-L1-L3 (Ensemble), and Model-Synthetic (Ensemble)) and four configurations for each model, this led to 4 × 4 = 16 configurations overall. These configurations allowed us to systematically analyze how different hyperparameters, resolutions of data, and model configurations influenced prediction accuracy and PPM error.

We extended our analysis by training a Decision Tree Regressor (using the squared-error criterion) and a Random Forest Regressor (50 estimators). For these models, we used mass and mobility features jointly as inputs. We formulated chemical formula assignment as a multi-output regression task in which each target vector contains the element counts for C, H, O, N, and S.

For each ion i, the input feature vector

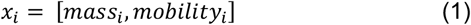

was mapped to a target vector of elemental counts

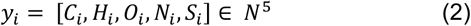

The models learns the mapping between input and output by minimizing the squared error loss across all elements

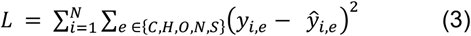

This framing helps to preserve the ordinal structure of elemental counts and avoids the combinatorial explosion that a multiclass classification formulation would introduce. This illustration is shown in Figure 1(b).

In our evaluation, both Matched Annotation (Formulas matched with rule-based tool) and New Annotations (Formulas with less than 1 PPM error and different from rule-based tool) are classified as True Annotations (TA), as Composer does not represent an absolute ground truth. We evaluated the model results using the assignment rate defined as:

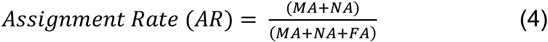

where MA represents matched snnotation, NA refers to new annotations, and FA denotes false annotations (with an error greater than 1 ppm). This metric measures the percentage of valid assignments, considering both:

1. Matched Annotations that match the Composer’s results, and
2. New Annotations that differ from the Composer’s assignments.

To evaluate the findings of DTR and RFR, we used two metrics. The first metric is the Formula-level Accuracy (FA) which are predictions that exactly match all true elemental counts and Element-level Accuracy (EA) which is the accuracy for each standalone element.

## 3 Results

For the Model-L1, both Euclidean and Manhattan distance metrics with K=1 and K=3 achieved identical outcomes with 3,202 matched annotations, 6 new annotations, and 840 false annotations (More than 1 PPM Error). It achieved an assignment rate of 79.27 %.

The Model-L3 showed a substantial improvement, achieving approximately 95% assignment rate. Specifically, for both K=1 and K=3, Euclidean and Manhattan distances performed nearly identically, with matched annotations around 3,831–3,835, false annotations between 202–205, and assignment rates of 94.94-95.01%. These results show the significance of higher mass resolution data.

For Model-L1-L3 (Ensemble), the model showed 3846-3851 matched annotations, 8-10 new annotations, and 188-191 false annotations. In this setting. K=1 has assigned more formulas than K=3, while the impact of distance metrics is identical.

The Model-Synthetic (Ensemble) achieved the highest performance, with an assignment rate of 99.90 %. For K=1 and K=3 under Euclidean and Manhattan metrics, matched annotations were 3934-3938, new annotations were 105–107, and false annotations were only 4-6. As this approach added synthetic data to the training data, it expanded the model coverage by increasing the number of data points. Model-Synthetic (Ensemble) space promotes the discovery of new valid formulas more than other approaches. These results are shown in Table 1.

**Table 1.**
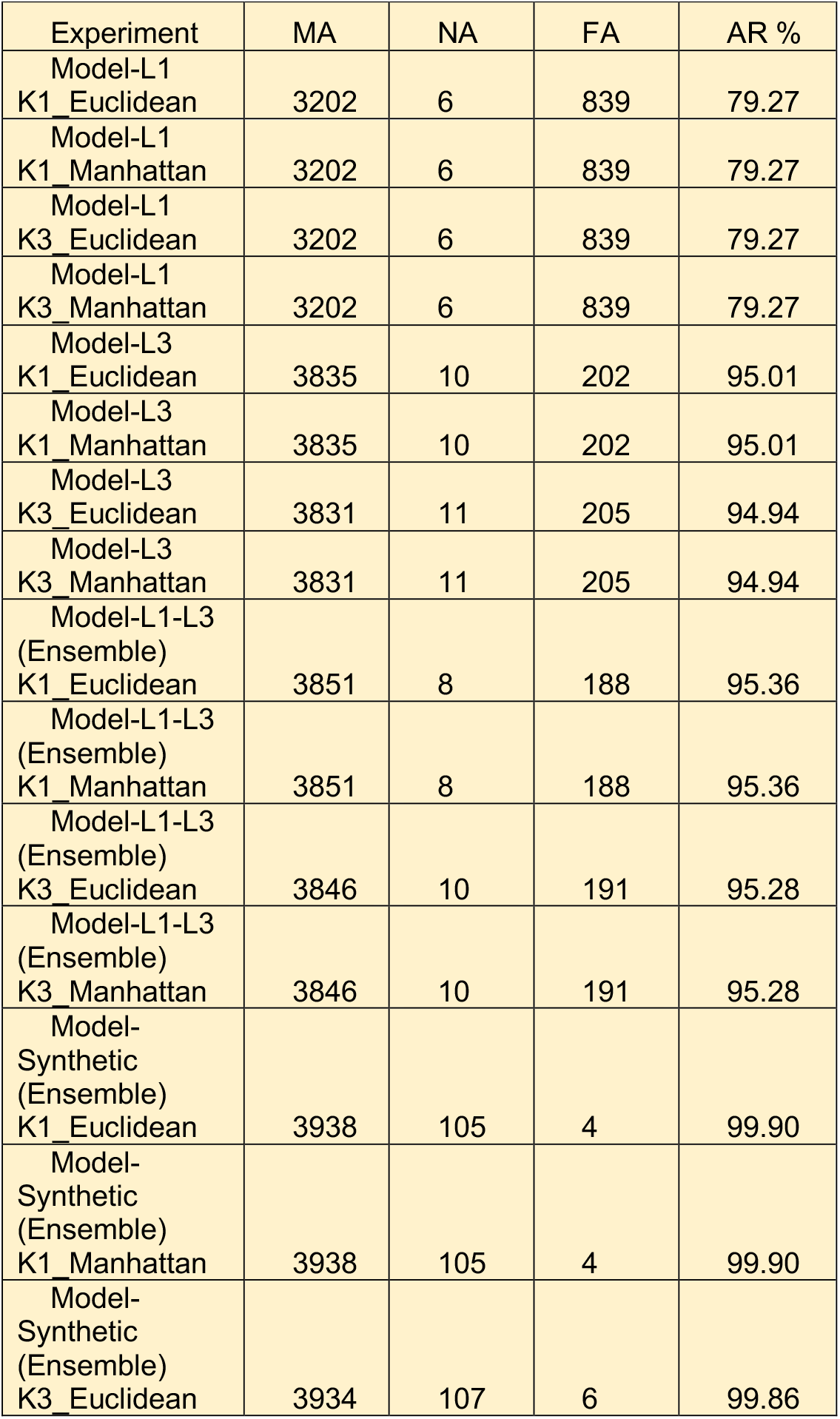
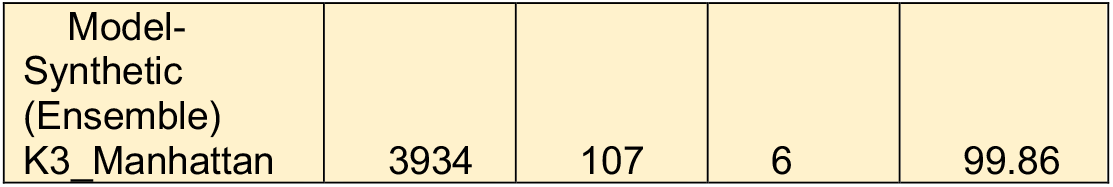
shows matched annotations (MA: number of assigned formulas same as rule-based tool), new annotations (NA: number of valid formulas(<1PPM) different from rule-based tool), false annotations (FA: number of formulas with more than 1 PPM error), and Assignment Rate (AR: Percentage of true annotations)

The confusion matrix analysis provides further insight into the classification performance across all experimental configurations. In this analysis, matched annotations and new annotations are considered true annotations, whereas predictions with more than 1 PPM error are considered as false annotations. For the Model-L1, both Euclidean and Manhattan metrics under K=1 and K=3 showed identical confusion matrix predictions, with a total of 3,208 true annotations and 839 false annotations. For Model-L3, the models showed significantly improved performance. For both distance measures and K values, the confusion matrix showed 3,842–3,845 true annotations and approximately 202–205 false annotations. The Model-L1-L3 (Ensemble) experiments predicted 3,856–3,859 true annotations and 188 to 191 false annotations. The Model-Synthetic (Ensemble) produced near-perfect confusion matrix outcomes. Across all configurations, a total of 4,041-4,043 true annotations and 4-6 false annotations were recorded. Overall, the confusion matrix evaluation confirms that all models achieved strong discriminative performance, with the synthetic dataset consistently producing the highest true annotations and only 4-6 false annotations, which demonstrates its robustness and generalization capability across different distance metrics and k parameters. The confusion matrix results are shown in Figure 2(a).

**Figure 2:**
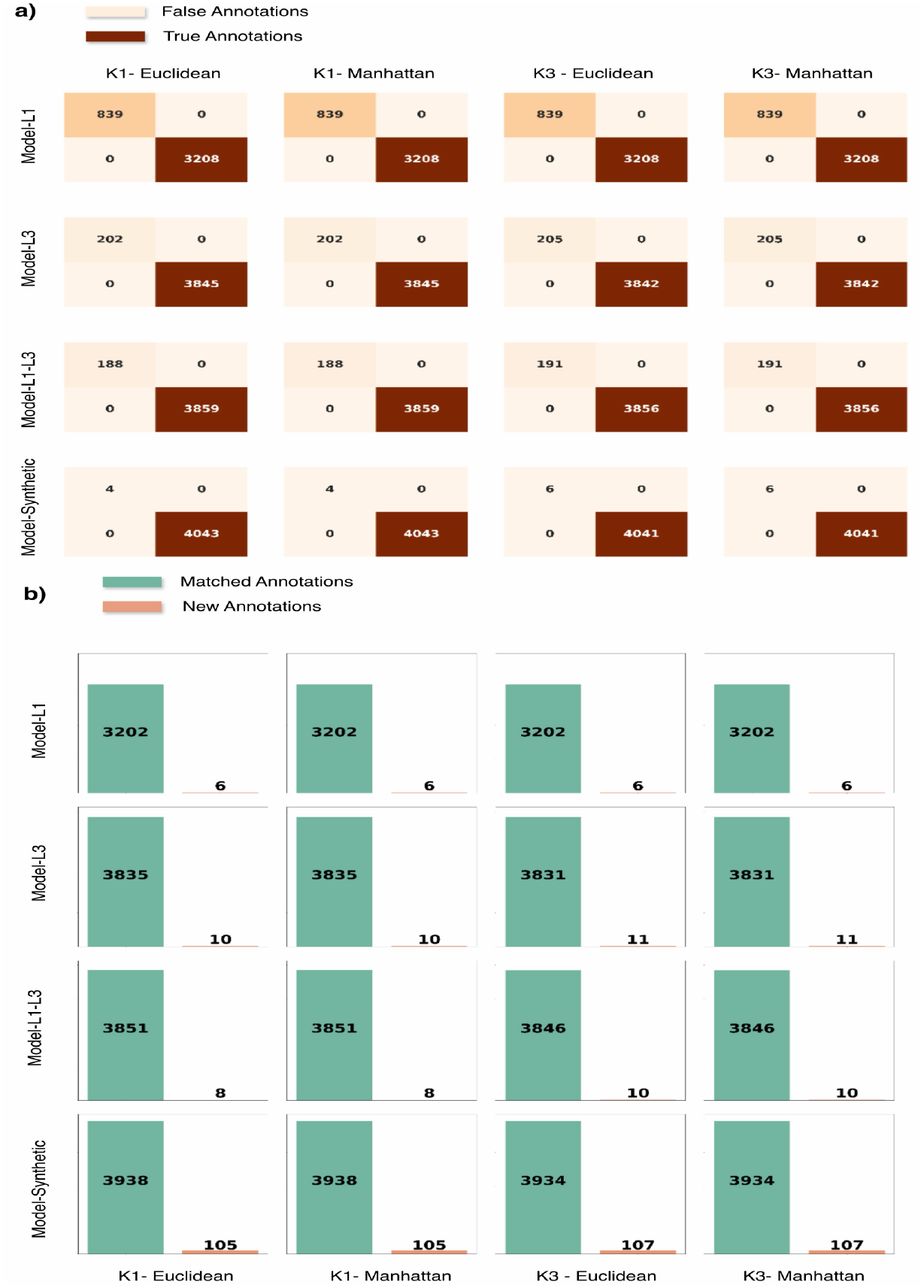
(a) Confusion Matrices showing true annotations of formulas for all configurations. Model-L1-L3 (Ensemble) and Model-Synthetic (Ensemble) approaches can annotate more formulas. (b) Bar Plots are showing number of matched annotations that match with the training set and number of new annotations assigned with less than 1 PPM error across all configurations.

To further examine the correctness and adaptability of the proposed approach, we performed an analysis focused on the matched annotations and new annotations. This highlights the model’s ability to accurately identify formulas that match with Composer and also annotate new formulas. The Model-L1 consistently annotated 3,202 formulas and generated 6 new annotations under both metrics for K=1 and K=3. Although the assignment rate remained moderate, these results demonstrate reliable performance on smaller-scale datasets. The Model-L3 predictive capability improved substantially across all metrics and K variations, it achieved between 3,831 and 3,835 matched annotations, along with 10 to 11 new annotations. This shows that the model generalizes effectively as the dataset complexity increases, with minimal redundancy between the known and newly discovered formulas. For Model-L1-L3 (Ensemble), the predictions showed improvement from Model-L1 and Model-L3. The approach has 3,846 to 3,851 matched annotations with 8 to 10 new annotations, indicating consistent performance and adaptability across data sources. The Model-Synthetic (Ensemble) provided more new annotations than any other approach. It shows the capacity for matched annotations as well as new annotations. It has 3934-3938 matched annotations, supplemented by 105 to 107 new annotations, demonstrating a strong ability to annotate new valid formulas. Overall, when focusing solely on the matched annotations and new annotations, the results reveal that the model not only achieves high reliability in recognizing existing patterns but also exhibits an impressive capacity to annotate new valid formulas, particularly evident in the synthetic dataset experiments. These results are shown in Fig.2(b).

Fig 3(a) illustrates the distribution of predicted mass-to-charge (m/z) errors (in parts per million, ppm) across all experimental configurations. For the L1, L3, and Model-L1-L3 (Ensemble), the error distributions are spread between 0 and 1 ppm, with a gradual decline in frequency as the error increases. This indicates that while most predictions are concentrated at low error values, a moderate number of cases exhibit higher deviations, suggesting a slight variability in model precision for these datasets. In contrast, the Model-Synthetic (Ensemble) exhibits a more skewed distribution, with the majority of predictions clustered below 0.5 ppm. This narrower spread demonstrates that the synthetic data achieves higher confidence in predictions across both Euclidean and Manhattan metrics, regardless of k size (K=1 or K=3). Overall, the figure confirms that the model maintains strong prediction confidence across all datasets, with the synthetic experiments providing the most confidence in predictions.

**Figure 3:**
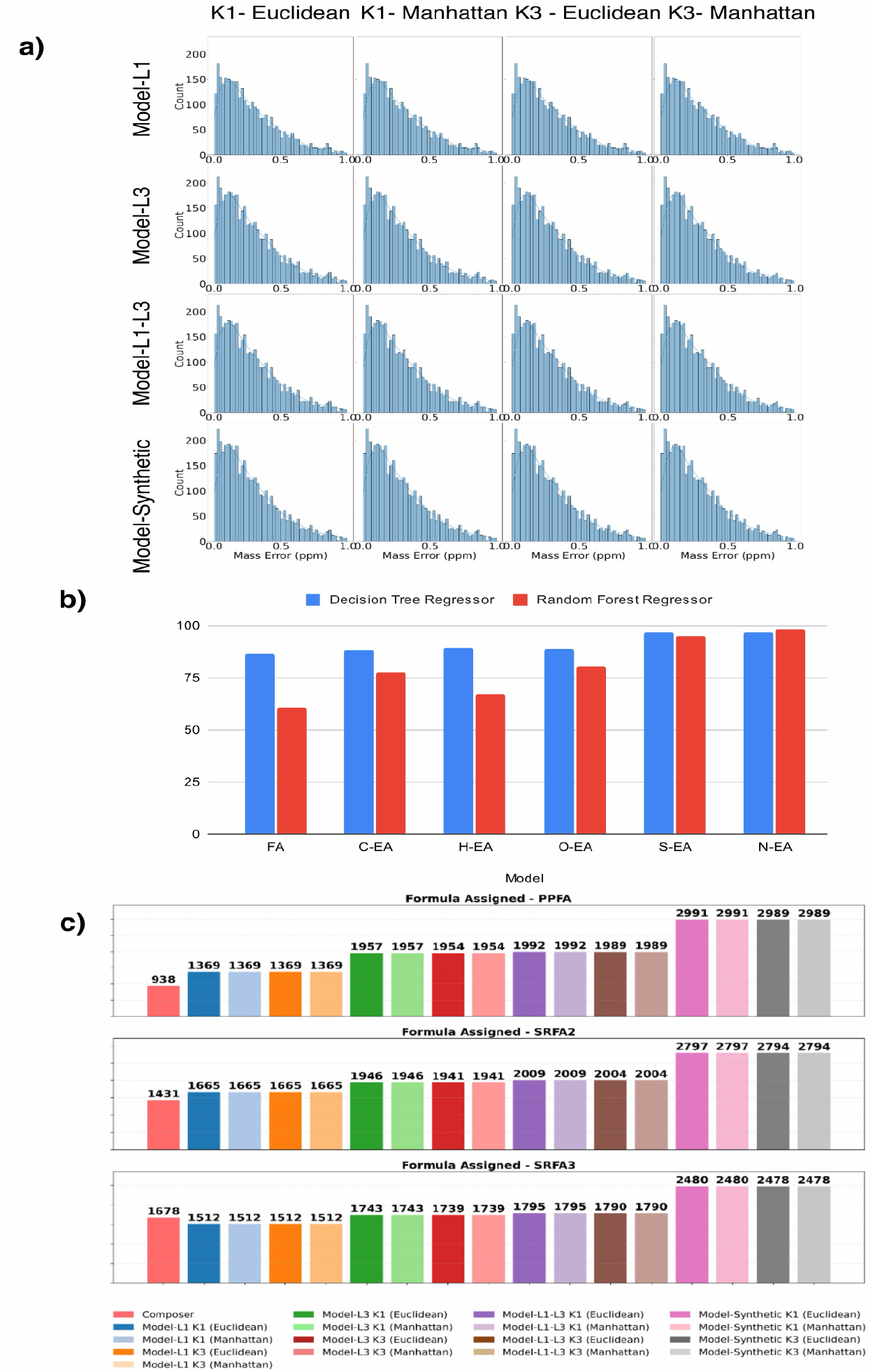
(a) Distribution of PPM errors for true annotations across all configurations. (b) Formula-level Accuracy (FA) and Element-level Accuracy (EA) for DTR and RFR (c) Comparison of Formula Assignments Peak lists, number of formulas annotated/assigned for PPFA, SRFA2 and SRFA3 with less than 1 PPM error.

For the Decision Tree Regressor (DTR) and Random Forest Regressor (RFR), we evaluated performance using both formula-level accuracy (FA) and element-level accuracy (EA). DTR achieves an FA of 86.5%, with element-level accuracies of 88.4% for C, 89.5% for H, 88.8% for O, and 96.6% for both S and N. In contrast, RFR attains an FA of 60.4%, with EA values of 77.6% for C, 67.1% for H, 80.4% for O, 94.9% for S, and 98.2% for N. These results are shown in Figure 3(b).

Figure 3(c) presents the distribution of molecular formula predictions for the PPFA, SRFA2, and SRFA3 peak lists. These peak lists do not have annotations (Formula annotated using rule-based tools), so all the predictions with mass error below 1 ppm are considered valid assignments. The goal here is to assign formulas to peaks. Each bar displays the number of annotations with mass errors below 1 ppm across all configurations. For PPFA, the annotations range from 1,369 to 2,991, reflecting a strong correlation between the observed and calculated masses across the model configurations. Similarly, SRFA2 shows a comparable distribution, with the number of formulas assigned extending from 1,665 to 2,797. SRFA3 shows formula assignments from 1,512 to 2,480. Composer assigned 938 formulas for PPFA, 1,431 for SRFA2, and 1,678 for SRFA3, while Model-Synthetic assigned 2,991 formulas for PPFA, 2,797 formulas for SRFA2, 2,480 formulas for SRFA3. In total, Composer annotated 4047 formulas, Model-L1-L3 annotated 5796 formulas and Model-Synthetic annotated 8268 Formulas. This shows 43% more formula assignment/annotation for Model-L1-L3 and 2x formula assignment/annotation for Model-Synthetic.

## 4 Conclusion

This paper presented a machine-learning approach for complex fulvic acid fraction (FA-DOM) chemical formula assignment from ultra-high-resolution mass spectrometry data. We generated a high-quality and ultra-high resolution data, including 1 PPM (L1), 0.2-0.4 PPM (L2) and 0.15 PPM (L3) mass resolutions, along with a large scale synthetic data of chemically plausible CHONS formulas. This dataset provide comprehensive coverage of experimental conditions to train and benchmark machine learning models. Our developed KNN pipeline was evaluated in a comprehensive manner across multiple datasets using Euclidean and Manhattan distance metrics under different neighborhood settings. KNN-Synthetic achieved 99% assignment rate and Model-L1-L3 annotated 43% more formulas than traditional rule-based tool. To further validate the robustness of our approach, we also benchmarked tree-based models, including Decision Tree Regressor (DTR) and Random Forest Regressor (RFR). DTR achieved 86.5% formula-level accuracy, while RFR reached 60.4%, with both models showing high element-level accuracy, particularly for elements such as S and N. These results confirm the reliability of the framework beyond a single model type and highlight its flexibility for different learning paradigms. Our experimental results (across 3 standards) demonstrate consistently high assignment rates across diverse datasets. Notably, the method achieved a strong performance on the synthetic data, where the assignment rate exceeded 99% and up to 2X formulas are annotated as compared to the existing state of the art methods. The mass error remained below 0.5 ppm for the majority of molecular formula predictions.

The Model-L1, Model-L3, and Model-L1-L3 (Ensemble) experiments also showed robust performance, with assignment rates ranging from 79% to 95%, confirming the reliability of the approach even under different mass resolutions. Analysis of the confusion matrix highlighted high true annotations and minimal false annotations, while the matched annotations and new annotations results underscored the model’s capability, not only to recognize chemical formulas that match with traditional tool but also to discover new formulas. Our findings validates the effectiveness and generalizability of the proposed approach for accurate molecular formula assignment. Future work includes extending the framework to larger and more diverse datasets, integrating new features, annotating/assigning more than one formula to one peak, and more advance machine learning approaches for complex DOM and meta-proteomics data sets.

## 5 Data Availability

All training and test datasets, and analysis results are publicly available at https://github.com/pcdslab/dom-formula-assignment-using-ml, including L1 and L3 FT-ICR MS training data, synthetic molecular formulas, and the L2 test dataset. You can also access them on hugging face https://huggingface.co/datasets/SaeedLab/dom-formula-assignment-data.

## 6 Code Availability

The complete pipeline implementation is available as open-source software at https://github.com/pcdslab/dom-formula-assignment-using-ml. Pre-trained models for all 16 configurations are hosted on Hugging Face Model Hub at https://huggingface.co/SaeedLab/dom-formula-assignment-using-knn.

## ACKNOWLEDGEMENTS

This research was supported by the NIGMS of the National Institutes of Health (NIH) under award number: R35GM153434 (FS). The authors were further supported by the National Science Foundations (NSF) under the award number: NSF OAC-2312599 (FS) and NSF CHEM 2304837 (FFL and FS). The content is solely the responsibility of the authors and does not necessarily represent the official views of the National Institutes of Health and/or National Science Foundation. The authors acknowledge the personnel of the Advance Mass Spectrometry Facility at Florida International University as well as David Stranz and Sierra Analytics, Inc. for their support with the data acquisition and interpretation.

## Appendix A Details about Formula Assignments

### A.1 Unique Formula Assignment for Peaklists

This shows the number of unique formulas annotated by each ML method. If more than 1 peak are assigned the same formulas, only 1 peak is selected. This is shown in Figure 4.

**Figure 4:**
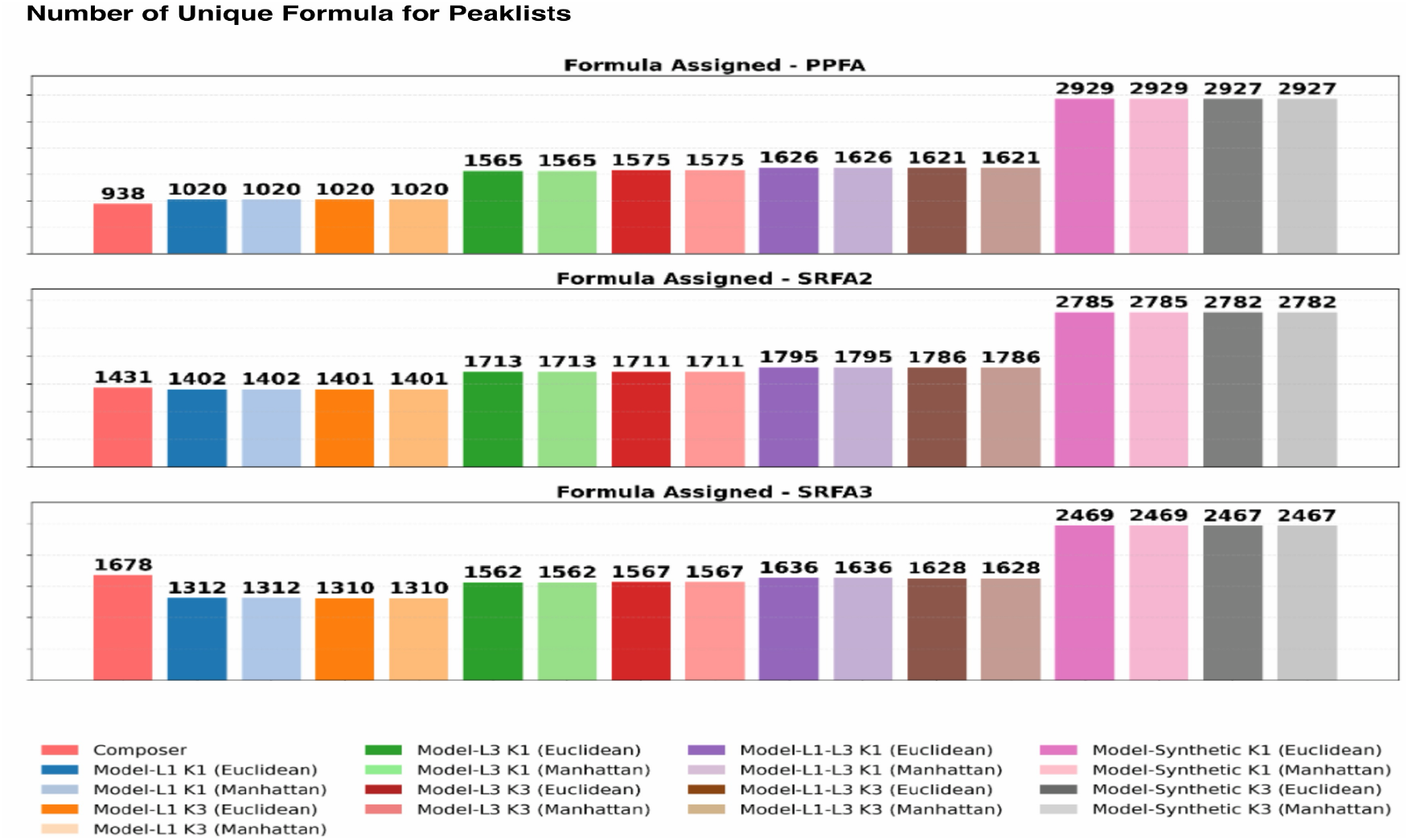
Unique formulas annotated from each model. If one more than one peaks annotated the same formula, only one peak is picked.

### A.2 Heteroatom distribution Plots

We show the heteroatom distribution of the plots. It shows the formulas assigned across different heteroatom classes. These results show that machine learning models assign the formulas in the same distribution as denovo tools. It is shown in Figure 5.

**Figure 5:**
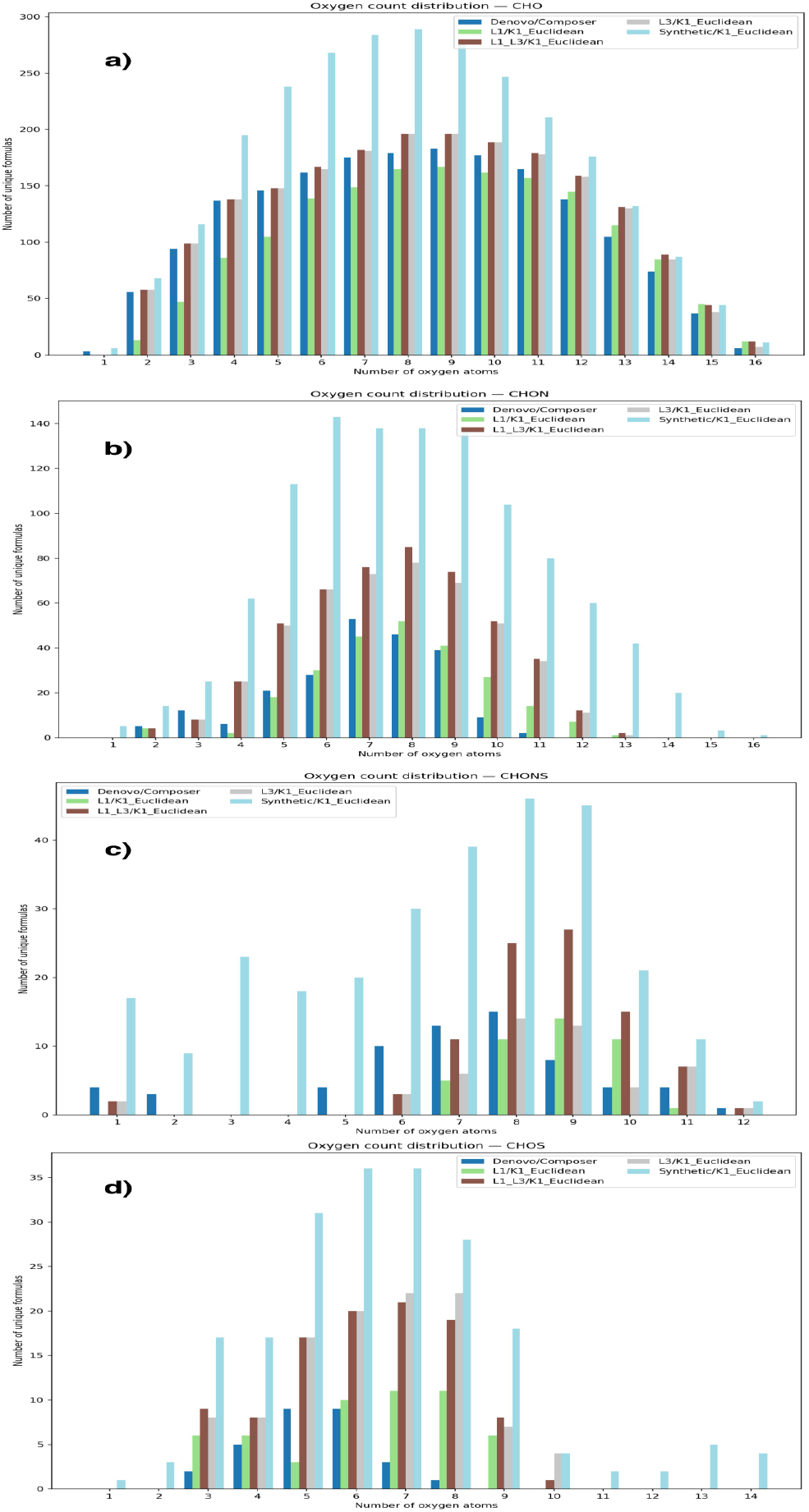
Distribution of formulas assigned/annotated for number of oxygen atoms across heteroatom classes a) CHO b) CHON c) CHONS and, d) CHOS.

